# Before trilobite legs: *Pygmaclypeatus daziensis* reconsidered and the ancestral appendicular organization of Cambrian artiopods

**DOI:** 10.1101/2021.08.18.456779

**Authors:** Michel Schmidt, Xianguang Hou, Dayou Zhai, Huijuan Mai, Jelena Belojević, Xiaohan Chen, Roland R. Melzer, Javier Ortega-Hernández, Yu Liu

**Affiliations:** MEC International Joint Laboratory for Palaeobiology and Palaeoenvironment, Yunnan University, 2 North Cuihu Road, Kunming 650091, People’s Republic of China; Yunnan Key Laboratory for Palaeobiology, Institute of Palaeontology, Yunnan University, North Cuihu Road 2, Kunming 650091, People’s Republic of China; Bavarian State Collection of Zoology, Bavarian Natural History Collections, Münchhausenstr. 21, 81247 München, Germany; Department Biology II, Ludwig-Maximilians-Universität München, 82152 Planegg-Martinsried, Germany; GeoBio-Center, Ludwig-Maximilians-Universität Munich, Luisenstr. 37, 80333 München, Germany; Museum of Comparative Zoology and Department of Organismic and Evolutionary Biology, Harvard University, 26 Oxford Street, 791 Cambridge, MA 02138, USA

**Keywords:** Cambrian, Chengjiang, heteronomy, computed tomography, exceptional preservation

## Abstract

The Cambrian Stage 3 Chengjiang biota in South China is one of the most influential Konservat-Lagerstätten worldwide thanks to the fossilization of diverse non-biomineralizing organisms through pyritization. Despite their contributions to understanding the evolution of early animals, several Chengjiang species remain poorly known due to their scarcity and/or incomplete preservation. Here, we use micro-computed tomography to reveal in detail the ventral appendage organization of the enigmatic non-trilobite artiopod *Pygmaclypeatus daziensis* – one of the rarest euarthropods in Chengjiang – and explore its functional ecology and broader evolutionary significance. *Pygmaclypeatus daziensis* possesses a set of uniramous antennae and 14 pairs of post-antennal biramous appendages, the latter of which show an unexpectedly high degree of heteronomy based on the localized differentiation of the protopodite, endopodite and exopodite along the antero-posterior body axis. The small body size (less than 2 cm), presence of delicate spinose endites, and well-developed exopodites with multiple paddle-shaped lamellae on the appendages of *P. daziensis* indicate a nekto-benthic mode of life, and a scavenging/detritus feeding strategy. *Pygmaclypeatus daziensis* shows that appendage heteronomy is phylogenetically widespread within Artiopoda – the megadiverse clade that includes trilobites and their relatives with non-biomineralizing exoskeletons – and suggests that a single exopodite lobe with paddle-like lamellae is ancestral for this clade.

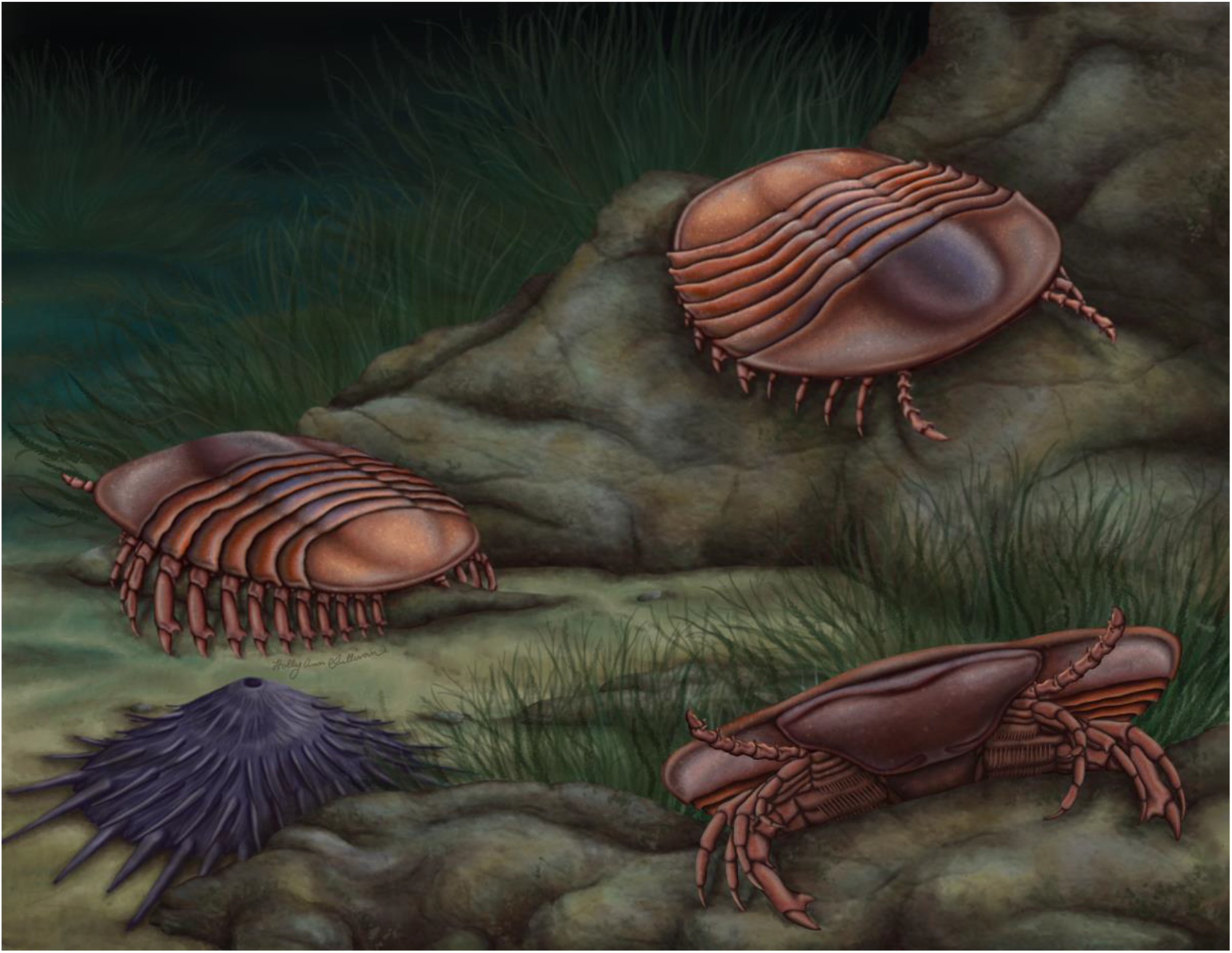

**Cover image:** Morphological reconstruction of the non-trilobite artiopod *Pygmaclypeatus daziensis* from the early Cambrian (Stage 3) Chengjiang biota in south China. Artwork by Holly Sullivan (https://www.sulscientific.com/).

## INTRODUCTION

The artiopods comprise a megadiverse clade of crown-group euarthropods with a distinctively flattened dorsal exoskeleton, and are among the dominant epibenthic animals in lower Paleozoic marine deposits (1, 2, 3, 4). Trilobites are by far the most ubiquitous members of Artiopoda thanks to the higher preservation potential granted by their calcitic dorsal exoskeleton (5), but the group also contains a significant diversity of non-biomineralizing taxa that are primarily known from Cambrian Konservat-Lagerstätten around the world, particularly from localities in South China (e.g. 6), North America (e.g. 7), North Greenland (e.g. 8, 9) and South Australia (e.g. 10, 11). Although these “soft-bodied” non-trilobite artiopods are moderately disparate in terms of the dorsal exoskeletal morphology, their appendicular organization has been traditionally regarded as a largely homonomous series of biramous gnathobasic limbs consisting of multi-segmented endopodites, and lamellate or broad flap-like exopodites (1, 06, 12, 13). Thus, most non-trilobite artiopods are usually considered as epibenthic deposit feeders or generalized scavengers/predators based on their conservative limb construction (e.g. 6, 14), with only rare cases of clear anatomical adaptations for more specialized dietary preferences such as durophagy (15, 16, 17).

The application of micro-computed tomography (micro-CT) to study pyritized macrofossils from the Cambrian (Stage 3) Chengjiang biota in South China can inform the ventral three-dimensional organization of fossil euarthropods in greater detail than would be possible under conventional imaging techniques (e.g. 18, 19, 20, 21). Recent studies of the appendicular organization in the xandarellid *Sinoburius lunaris* (22) and the nektaspid *Naraoia spinosa* (23) demonstrate a higher degree of morphological differentiation in artiopods than previously appreciated (e.g. 01, 06, 07, 24), including the regionalization of the biramous appendages along the antero-posterior body axis, as well as functional changes in feeding strategy during ontogeny. Although these examples demonstrate the cryptic complexity of the appendages in two of the main lineages of Trilobitomorpha (25), namely the group that includes all those forms more closely related to trilobites than to vicissicaudates (xenopods, cheloniellids aglaspidids; see 3), the organization of early branching members of Artiopoda remains enigmatic due to the scarcity of fossil taxa with well-preserved and clearly exposed limbs.

In this study, we describe the appendicular organization of the rare and enigmatic non-trilobite artiopod *Pygmaclypeatus daziensis* (26) and explore its ecological and evolutionary implications. More than 20 years after its original description, the morphology, ecology and affinities of *P. daziensis* remain problematic (27, 28). Type material from Chengjiang hints at the preservation of appendages (Fig. 1a), but fine morphological details concealed within the rock matrix are inaccessible through conventional photography and preparation methods. The early branching phylogenetic position of *P. daziensis* within the artiopod tree (10, 29) makes it directly significant for reconstructing the ancestral appendage organization and functional morphology of this major clade of Paleozoic euarthropods.

**Figure 1.**
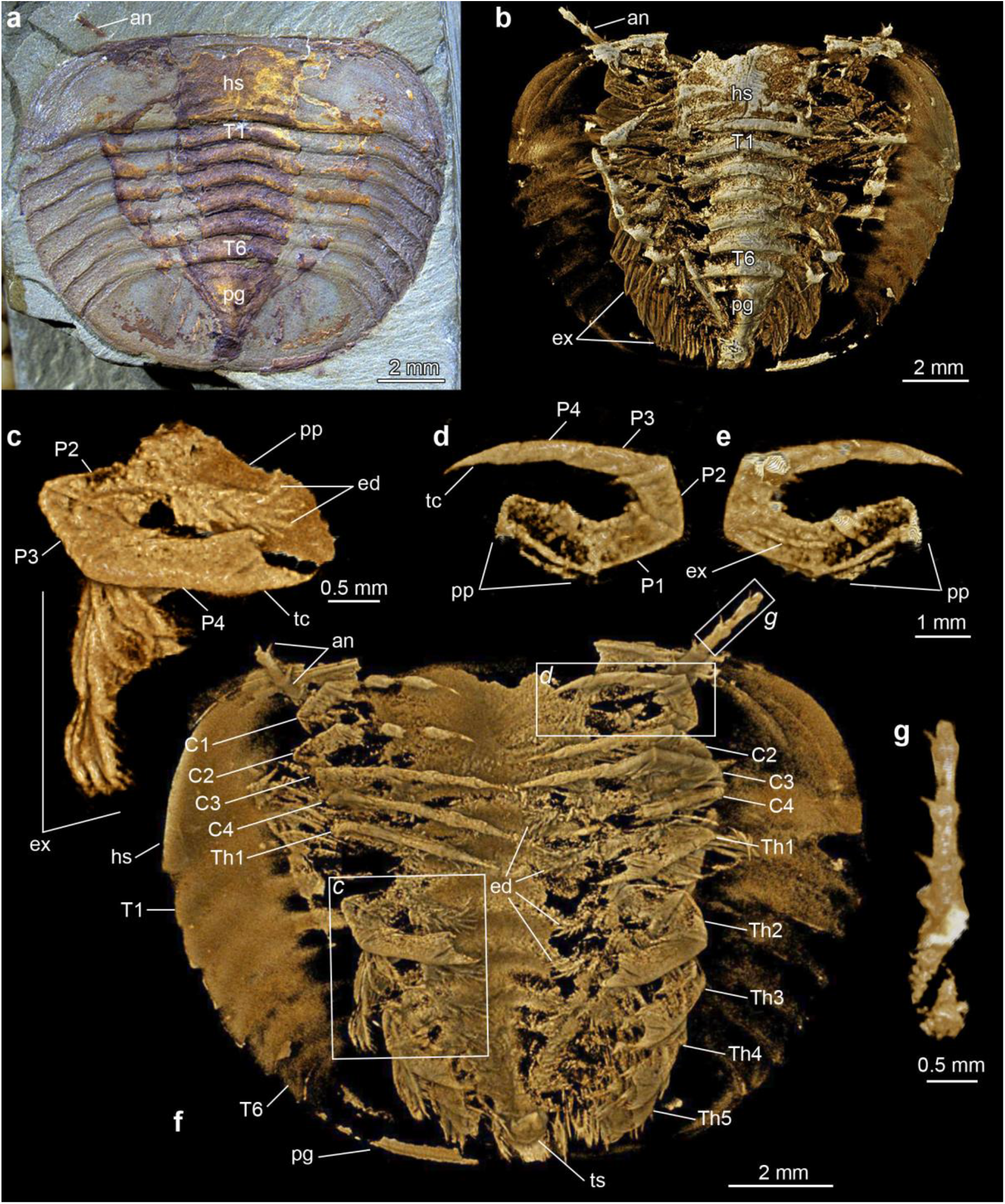
*Pygmaclypeatus daziensis* from the Cambrian (Stage 3) Chengjiang biota of South China. **a**. Specimen YKLP 11427, photographed under light microscopy. **b**. Dorsal view of three-dimensional computer model based on X-ray tomographic data rendered in Drishti [32]. **c**. Isolated three-dimensional model of second thoracic appendage in ventral view. **d**. Isolated three-dimensional model of first post-antennal cephalic appendage in ventral view. **e**. Isolated three-dimensional model of first post-antennal cephalic appendage in dorsal view, showing attachment site of elongate stenopodous exopodite. **f**. Ventral view of three-dimensional computer model based on X-ray tomographic data, exceptionally preserved appendage morphology. **g**. Isolated three-dimensional model of left antenna, showing segmental boundaries and accessory spines. Abbreviations: *an*, antenna; *Cn*, cephalic post-antennal appendage pair number; *ed*, enditic spine bristles; *ex*, exopodite; *hs*, head shield; *pp*, protopodite; *Pn*, podomere number; *tc*, terminal claw; *Tn*, tergite number; *Thn*, thoracic appendage pair number; *ts*, tailspine.

## MATERIALS AND METHODS

### Materials

The studied material includes four specimens (YKLP 11427, 11428, 13928, and 13929a of *Pygmaclypeatus daziensis* collected from the Yu’anshan Member of the Chiungchussu Formation, Dazi section of Haikou in Yunnan Province, South China. All specimens are deposited at the Yunnan Key Laboratory for Palaeobiology at Yunnan University, Kunming. All specimens are preserved in dorsal view, and are replicated in pyrite and/or iron oxides as typically observed in fossil euarthropods from the Chengjiang Lagerstätte (30, 31).

### Fossil imaging

Light photography was performed with either a Canon EOS 5DSR camera (DS126611) with a MP-E 65mm macro photo lens or a Keyence VHX-5000 digital microscope. Except for YKLP 13929, scans of all the other specimens were performed using an Xradia 520 Versa (Carl Zeiss X-ray Microscopy, Inc., Pleasanton, USA) either at the Yunnan Key Laboratory for Palaeobiology (YKLP 11428) or at the Institute of Geology and Geophysics, Chinese Academy of Sciences (YKLP 11427, 13928). YKLP 13929 was scanned using a Phoenix Nanotom (GE Sensing & Inspection Technologies) cone-beam CT scanner located at the Bavarian State Collection of Zoology, München. Scanning parameters are as the following: YKLP 11427 (Fig. 1b-g): Beam strength: 60kV/5w, Filter: no, Resolution: 8.79µm, Number of TIFF images: 2030; YKLP 11428 (Fig. 2): Beam strength: 50kV/4w, Filter: no, Resolution: 8.74µm, Number of TIFF images: 1937; YKLP 13928 (Extended Data Figure 1c, d): Beam strength: 60kV/5w, Filter: no, Resolution: 8.79µm, Number of TIFF images: 2030; YKLP 13929 (Extended Data Figure 1a, b): Beam strength: 130kV/14w, Filter: no, Resolution: 17.5µm, Number of TIFF images: 2397. Volume and surface renderings for YKLP 11428, 11429 were produced in Drishti 2.4 (32). Three-dimensional morphological reconstructions were created in Blender 2.9 (33).

**Figure 2.**
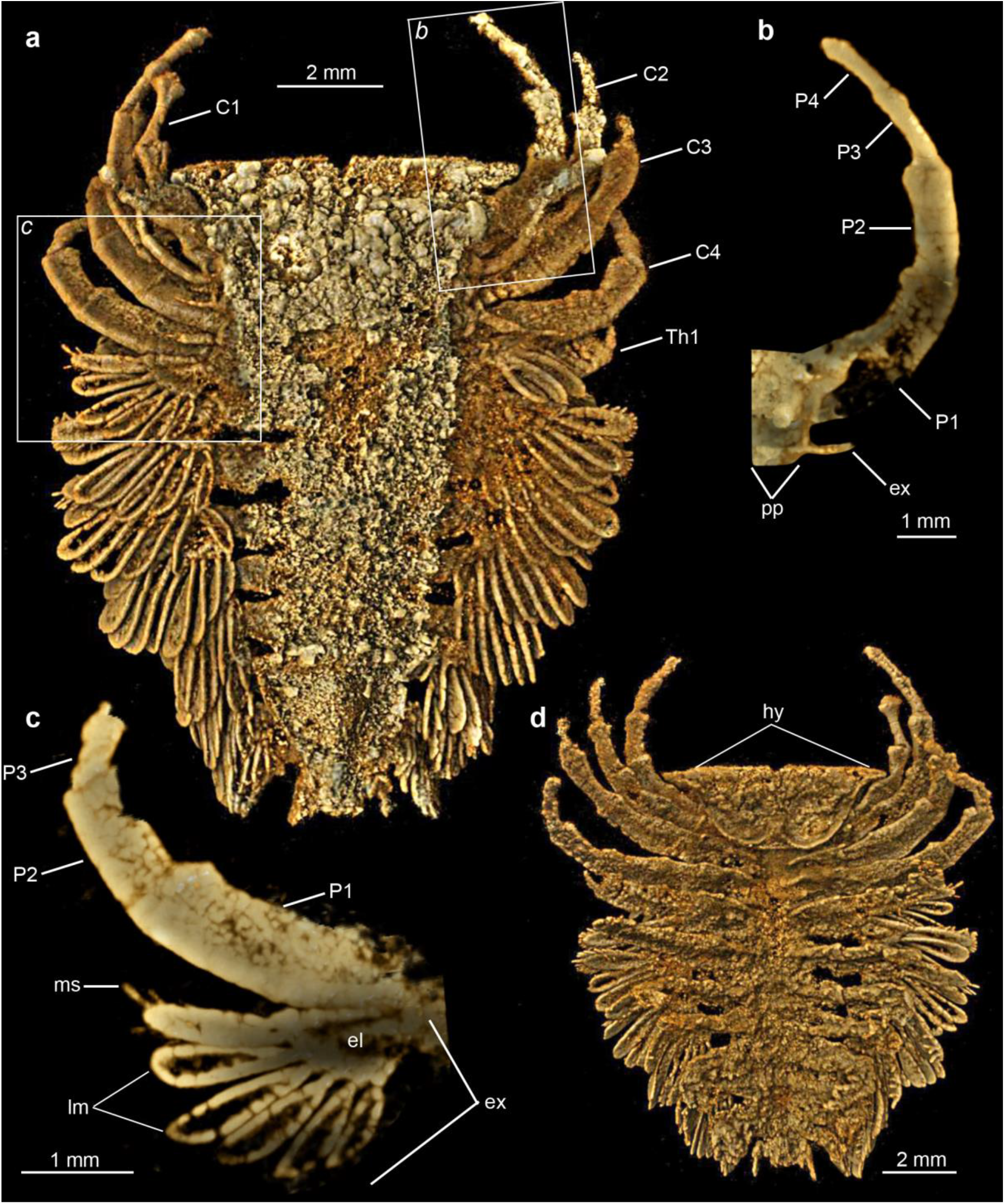
*Pygmaclypeatus daziensis* from the Cambrian (Stage 3) Chengjiang biota of South China. **a**. Specimen YKLP 11428, dorsal view of three-dimensional computer model based on X-ray tomographic data rendered in Drishti (32); dorsal exoskeleton tergites removed to illustrate the appendage organization. **b**. Isolated three-dimensional model of first post-antennal cephalic appendage in dorsal view, showing attachment site of elongate stenopodous exopodite. **c**. Isolated three-dimensional model of first thoracic appendage in dorsal view, showing morphology and attachment site of well-developed exopodite with paddle-shaped lamellae; note that marginal setae are only present on the distal-most lamella. **d**. Ventral view of three-dimensional computer model based on X-ray tomographic data, showing well-preserved hypostome. Abbreviations: *Cn*, cephalic post-antennal appendage pair number; *el*, exopodite lobe; *ex*, exopodite; *hy*, hypostome; *lm*, paddle-shaped lamellae; *ms* marginal setae; *pp*, protopodite; *Pn*, podomere number; *Thn*, thoracic appendage pair number.

### Phylogenetic analysis

The character matrix used for the phylogenetic analysis consists of 65 taxa and 92 characters. We employed an updated version of the dataset used by Chen et al. (22), including new morphological characters and updates to previous codings (Supplementary Information). Parsimony analyses were performed with TNT V1.5 (34) under New Technology Search, using Driven Search and Sectorial Search, Ratchet, Drift, and Tree fusing options activated with standard settings. The analysis was set to find the minimum tree length 100 times and to collapse trees after each search. All characters were treated as unordered. Analyses were performed under equal and implied weights.

## RESULTS

### Systematic Paleontology

Euarthropoda Lankester, 1904 (35)

Artiopoda Hou and Bergström, 1997 (06)

*Pygmaclypeatus* Zhang, Han, and Shu 2000 (26)

### Emended diagnosis

Small artiopod with dorsoventrally flattened exoskeleton, broader than long. Dorsal exoskeleton with poorly defined axial region without clearly developed axial furrows. Cephalon short, covering widely conterminant hypostome, uniramous antennae, and four appendage pairs. Trunk consists of six freely articulating tergites, each covering a single appendage pair. Pygidium nearly isopygous, covering four appendage pairs, and short multiarticulated tailspine. All post-antennal appendages biramous, consisting of protopodite, endopodite with five podomeres, and exopodite. Biramous appendage construction variable throughout the body; anterior three pairs of cephalic limbs lacking protopodal gnathobases and with reduced stenopodous exopodite, whereas all remaining posterior pairs have delicate spinose endites on protopodite, and well-developed exopodite lobe bearing thick paddle-shaped lamellae. Modified from Zhang et al. (26).

### Remarks

*Pygmaclypeatus* is among the rarest fossil taxa in the Chengjiang biota, with an estimated total of around 10 specimens available to date (28). Since its original description (26), only one study has made further contributions to the morphology and preservation of *Pygmaclypeatus*, namely the recognition of a short multiarticulated tailspine (27). The phylogenetic position of *Pygmaclypeatus* within Artiopoda has also received little attention, although most studies concur in close affinities with the similarly enigmatic *Retifacies* based on the presence of a pygidium with a multiarticulated tailspine (06, 10, 11), and *Squamacula* from Chengjiang based on their overall shield-like shape and body proportions (36). Our new morphological data allows us to produce a more accurate diagnosis for *Pygmaclypeatus* in the context of Artiopoda.

### Pygmaclypeatus daziensis Zhang, Han and Shu, 2000

2000 Zhang, Han, and Shu, pp. 980, Fig. 1 (26)

2004 Xu, pp. 331, Plate 1, Figs 4, 5 (27)

**Figure 3.**
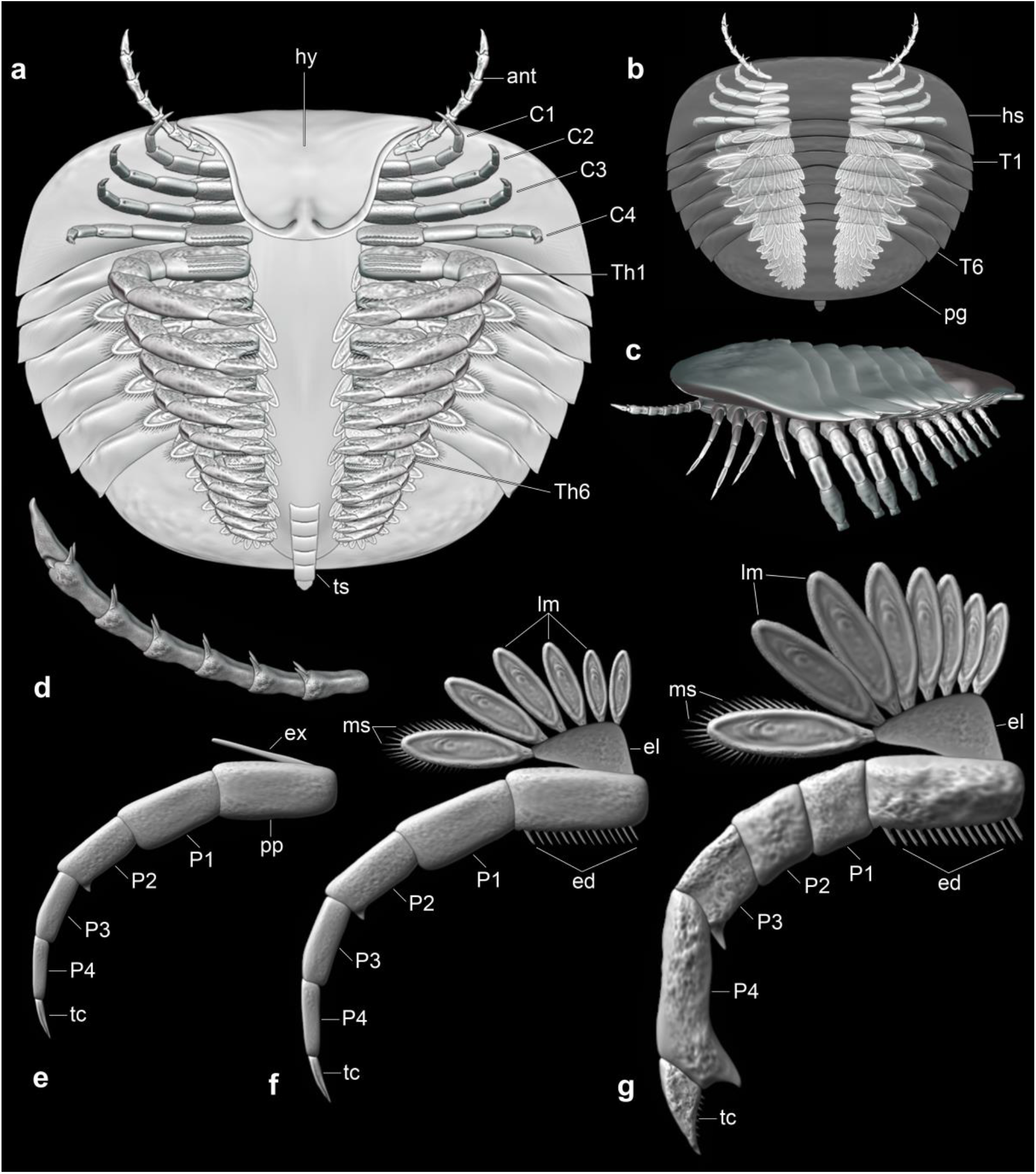
Three-dimensional morphological reconstruction of *Pygmaclypeatus daziensis*. **a**. Ventral view. **b**. Dorsal view, with transparent dorsal exoskeleton to show organization of appendages. **c**. Lateral view showing habits of ventral appendages. **d**. Antennae. **e**. Morphology of first to third post-antennal cephalic appendage pairs. **f**. Morphology of fourth post-antennal cephalic appendage pair. **g**. Morphology of thoracic and pygidial appendage pairs. Abbreviations: *an*, antennae; *Cn*, cephalic post-antennal appendage pair number; *ed*, enditic spine bristles; *el*, exopodite lobe; *ex*, exopodite; *hy*, hypostome; *lm*, paddle-shaped lamellae; *ms* marginal setae; *pp*, protopodite; *Pn*, podomere number; *tc*, terminal claw; *Tn*, tergite number; *Thn*, thoracic appendage pair number; *ts*, tailspine.

**Figure 4.**
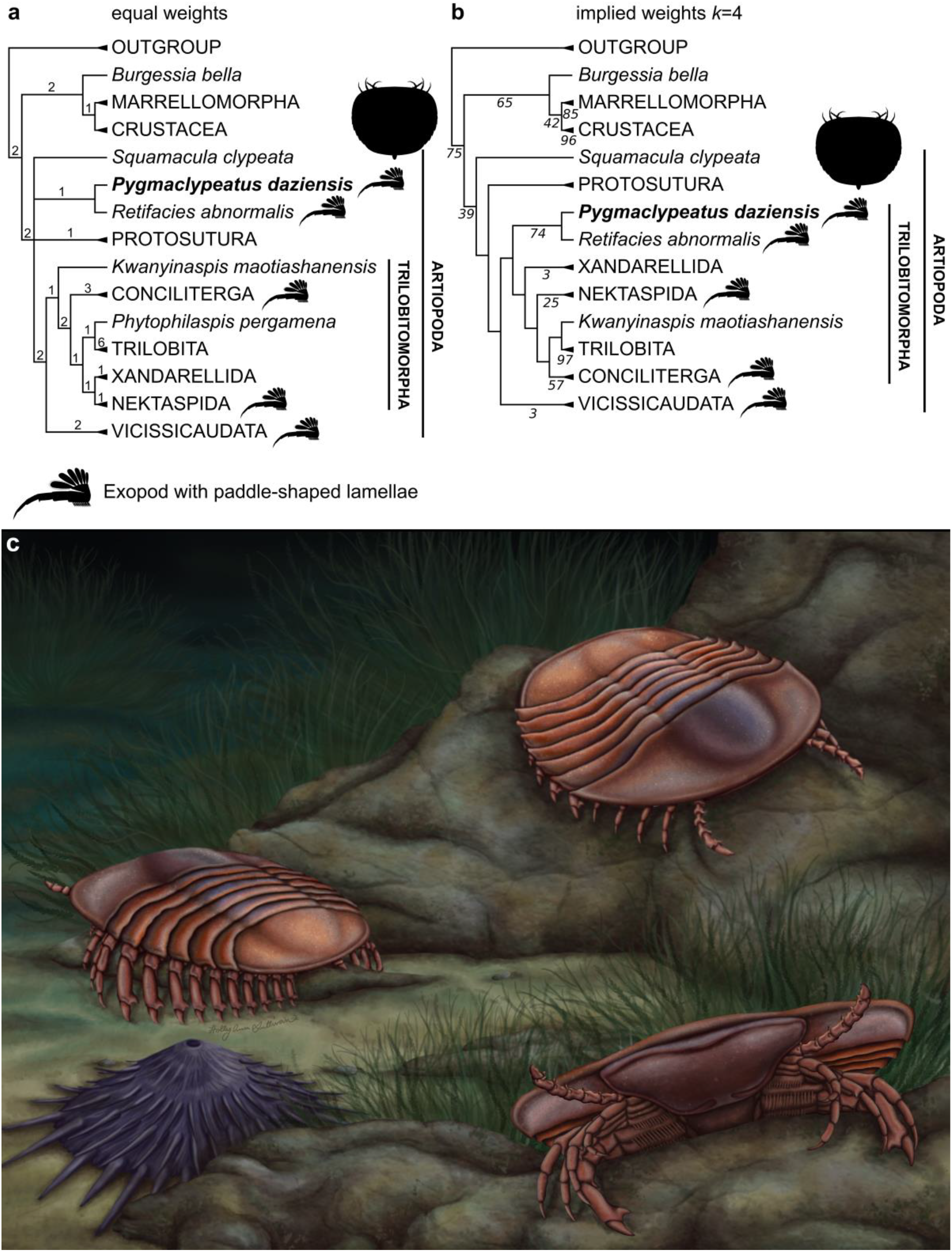
Results of parsimony-based phylogenetic analyses and morphological reconstruction. **a**. Strict consensus of four most parsimonious trees (265 steps; CI=0.419; RI=0.74) under equal weights; Bremer support values above nodes. **b**. Strict consensus of two most parsimonious trees (CI= 0.41; RI=0.73) under implied weights (*k=*4); nodal support expressed as symmetric resampling after 100 repetitions. Note the widespread occurrence of exopodites with paddle-shaped lamellae among trilobitomorphs (see text for discussion). **c**. Morphological reconstruction of *Pygmaclypeatus daziensis*. Artwork by Holly Sullivan (https://www.sulscientific.com/).

2017 Hou et al., pp. 197, Fig. 20.28 (28)

### Diagnosis

As for genus.

### Description

All known individuals are preserved in a dorsoventrally flattened position. Completely articulated specimens have a broad dorsal exoskeleton that is slightly wider than long, reaching up to 14 mm in length (sagittal), and a maximum width (transverse) of up to 17.5 mm measured at the level of the anterior trunk (Fig. 1). The dorsal exoskeleton has a shield-like rounded outline, with the anterior end of the body being easily distinguished by featuring a straight margin. The body consists of a cephalon, a trunk with six freely articulating tergites, and a pygidial shield associated with a multisegmented tailspine (Fig. 1, 2, 3). The cephalon comprises approximately 25% of the total body length (sag.). The cephalic outline is subrectangular with a straight anterior margin, rounded anterolateral edges that develop into posterior-facing acute genal angles, and a posterior margin that is gently reflexed adaxially (Fig. 1a). The dorsal surface of the cephalon is mostly featureless, without dorsal ecdysial sutures, and lacks evidence of eyes or other ocular-derived structures (*contra* 27). Although the main body axis is demarcated by a difference in coloration relative to the lateral areas of the cephalon, there is no indication of an elevated axial region or furrows. Previous studies have interpreted the presence of a rostral plate or anterior sclerite in the type material of *Pygmaclypeatus daziensis* (e.g. 10, 26, 37, 38). Close examination of our material, including micro-CT data, does not demonstrate the presence of a discrete anterior sclerite. Instead, the anterior margin of the head shield frequently features a transverse narrow band that appears to be a compaction artifact produced by the direct contact between the head shield and the ventral hypostome (Fig. 2d). We find no evidence of an anterior sclerite in *P. daziensis*.

The ventral side of the cephalon includes a hypostome, a pair of uniramous antennae, and four pairs of post-antennal biramous appendages (Fig. 1d, f; 2a, d). The hypostome is conterminant, directly attached to the anterior margin of the cephalon. The hypostome transversely covers ca. 33% of the width of the cephalon, and extends posteriorly to ca. 50% of the length of the cephalon, completely covering the proximal bases of two post-antennal appendages. Morphologically, the hypostome has a rounded posterior margin, including a small median notch that most likely indicates the position of the posterior-facing mouth as typically located in artiopods (39). The edges of the hypostome are strengthened by convex crescent-shaped ridges that meet adaxially, defining the median notch, conveying a bi-lobed appearance. The antennae are antero-laterally oriented, and attach ventrally in the cephalon close to the antero-lateral edges of the hypostome (Fig. 1f; 2d). The proximal portions of the antennae are well preserved in YKLP 11427 (Fig. 1b, f), consisting of at least four articles. The antennae appear to have a sub-cylindrical cross-section, and gently taper in width distally. The articles telescope into each other, with each featuring a pair of short spines located on their anterior margin, and facing adaxially relative to the main body axis. The full extent of the antennae is unknown due to their fragmentary preservation.

The first four post-antennal biramous appendages become progressively larger towards the posterior end of the cephalon, and include two morphological variants. The first three post-antennal appendages consist of a subrectangular protopodite attached to an endopodite with five podomeres, and a reduced stenopodous exopodite (Fig. 1d, e; 2b). The exopodites lack a clear indication of segmental boundaries, but it is unclear if this results from a preservation artifact or accurately reflects the original morphology. The endopodites are preserved in a strongly curved orientation facing towards the main body axis, and taper to an acute terminal claw. The podomeres are laterally compressed. In these limbs, there is little evidence of well-developed endites or any other processes on the protopodite or the endopodites in either of the examined specimens (Fig. 1d, e, f), suggesting that this absence is legitimate rather than taphonomic. The only exception is the presence of a short ventral spine on the anterior margin of the second endopodite podomere (Fig. 1d, 3e). The cephalic endopodites are prominent in size, and are the only appendages that extend beyond the exoskeleton margins (Fig. 2a). The exopodite is attached dorsally to the protopodite (Fig. 1d, 3e), and consists of a slender rod-shaped branch less than half the length of each corresponding endopodite. The fourth post-antennal biramous appendage pair has a similar – albeit more robust – overall endopodite organization, but differs substantially in the morphology of other components. The protopodite bears gnathobases along the ventral margin consisting of two or three densely populated rows of delicate spinose endites (Fig. 1f). The exopodite is fully developed, consisting of a subtriangular proximal base attached to the protopodite, and which bears up to six thick paddle-shaped lamellae (Fig. 2c). Each of the paddle-shaped lamellae shows a preferential preservation along the edges, suggesting that this region had strengthened cuticle relative to the interior face of the paddle. Within each exopodite, only the most distal paddle-shaped lamella bears a series of short marginal setae (Fig. 2c); the fact that this pattern is observed in several appendage pairs argues that this organization is not the result of a preservation artifact, and instead suggests that it is a legitimate biological feature. The variable orientation in the preservation of the paddle-shaped lamellae, as seen from dorsal view, indicates that these structures were freely articulated and allowed for some degree of motion through pivoting relative to the exopodite lobe (Fig. 2a, d).

The trunk of *P. daziensis* consists of six freely articulating tergites that overlap widely with each other (Fig. 1a, b). The trunk comprises approximately 50% of the total body length (sag.). All trunk tergites are much wider (trans.) than long (sag.), and share the same overall morphology. The axial region of the trunk suggests a weak elevation, which is also indicated by the anteriorly reflexed posterior margin of the tergites. The tergites have a sub-rectangular outline adaxially with straight margins, but become progressively curved posteriorly and abaxially, terminating in acute pleural angles. The first trunk tergite is approximately the same width (trans.) as the head shield, whereas the subsequent tergites gently taper in width towards the posterior end, conveying a rounded body outline (Fig. 1a, b). Each trunk tergite is associated with a single pair of biramous appendages on the ventral side. The trunk limbs are morphologically similar to the fourth post-antennal appendage pair, including the presence of a protopodite with delicate spinose ventral endites, endopodites with five podomeres, and exopodites with a triangular base that supports paddle-shaped lamellae with marginal setae (Fig. 1c). However, the trunk appendages differ in some details. The trunk endopodites are more robust and the podomeres are laterally compressed. Most significantly, the distalmost part of the endopodite, consisting of second to last podomere and the terminal claw, is modified into a strong sub-chelate termination (Fig. 1c). The terminal claw is broad and subtriangular, and features a wide area of articulation with the neighboring podomere that suggests a function as a moveable finger with a finely serrated inner edge, whereas the pre-terminal podomere is enlarged distally forming a robust thumb-like projection (fixed finger). Lastly, the larger size of the trunk exopodites can accommodate up to nine thick lamellae. A dorsal view of the articulated trunk appendage series shows that the ventral limbs are entirely concealed within the margins of the exoskeleton (Fig. 1a, b), which explains the difficulty of resolving these structures without the aid of micro-CT imaging.

The pygidium of *Pygmaclypeatus daziensis* consists of an unsegmented shield covering the posterior segments, and has a gently rounded posterior margin (Fig. 1a, b). The pygidium corresponds to approximately 25% of the total body length (sag.), but is narrower (trans.) than the cephalon and the anterior portion of the trunk. The weakly defined axial region of the cephalon and trunk continues into the pygidium, but ends at approximately the middle portion of the posterior shield with a rounded subtriangular termination. The ventral side of the pygidium shows the presence of at least four pairs of biramous appendages; these indicate that *P. daziensis* possess ca. 14 pairs of post-antennal appendage pairs (Fig. 1f; 2a, d). The morphology of the biramous appendages under the pygidium is identical to those in the trunk region, although the limbs become progressively smaller posteriorly and closer towards the midline. The posterior end of the body bears a short multiarticulated tailspine tucked underneath the body in YKLP 11427 (Fig. 1f), which corroborates the most recent revision of *P. daziensis* (27).

## DISCUSSION

### *Appendicular heteronomy in* Pygmaclypeatus

The use of micro-CT in pyritized Chengjiang fossils reveals the overall appendage morphology of *Pygmaclypeatus daziensis* in substantial detail (Fig. 1-3), made all the more striking by the fact that the known ventral organization of this taxon until now consisted of faint impressions of the hypostome and fragmentary antennae [26, 27, 28]. The appendages of *P. daziensis* combine a unique suite of ancestral and derived characters within the broader context of Artiopoda, and significantly, demonstrate a higher degree of appendage heteronomy than any other representative of this clade described to date.

Flagelliform antennae are widespread among artiopods, present in both trilobites (e.g. 13, 40) and their non-biomineralized relatives (e.g. 06, 23, 37); the Chengjiang xandarellid *Sinoburius lunaris* represents the only known case to date where the antennae are highly reduced and non-flagelliform (22). Paired spines on the uniramous antennae are also expressed in several non-biomineralizing artiopods such as *Retifacies, Kuamaia* and *Emeraldella* (e.g. 06, 37, 41), and even trilobites as in *Hongshiyanaspis* (13). Although the antennae of *P. daziensis* fall within the known diversity observed in artiopods, this is not the case for the post-antennal appendages. The organization of the first to third post-antennal biramous appendage pairs of *P. daziensis* is uncommon in the absence of well-developed spinose endites along the ventral edge, typical in most artiopods, with the exception of short spines associated with the anterior podomere margins in the endopodite (Fig. 1d, e; 2b). Intriguingly, this organization parallels that of the prosomal endopodites in the extant horseshoe crab (Xiphosura) *Limulus polyphemus* (42), both in terms of the number of podomeres and the presence of sparse short spines on their anterior edge. The occurrence of highly reduced stenopodous exopodites associated with the first to third post-antennal biramous appendages is also notable, as this precise morphology is unknown from most other artiopods. Comparable structures among Cambrian representatives include the flagelliform cephalic exopodites of *S. lunaris* (22) and *Sanctacaris uncata* (43), although these differ in being longer and multiarticulated, and the so-called proximolateral process found on the protopodite of *Sidneyia inexpectans* (02, 17, 44). Significantly, the latter comparison invites further parallels with the “flabellum” of *L. polyphemus*, a small lobe-like structure found on the protopodite of the sixth prosomal limb pair that has been interpreted as either a vestigial exopodite (45), or possibly an exite (46, 47).

The fourth pair of biramous appendages in *P. daziensis* has a more conventional organization based on the presence of prominent endites on the protopodite and a well-developed exopodite, but their precise morphology also has some uncommon features. The protopodite endites of *P. daziensis* are delicate and spinose, similar to those recently described in the protopodite of adult *Naraoia spinosa* from Chengjiang (23), and of the anterior walking legs of *Limulus polyphemus* (15, 17). However, the detailed morphology of the exopodite differs drastically from the elongate lobed organization observed in most artiopods, in which the exopodite typically consists of a rod-like shaft (e.g. *Misszhouia, Naraoia compacta, Xandarella*), or is differentiated into a slender proximal and large tear-shaped distal lobe (e.g. *Naraoia spinosa, Olenoides, Saperion*) (see fig. 4 in ref. 03). The fully developed exopodite of *P. daziensis* most closely resembles that of *Retifacies abnormalis* in the presence of a single robust lobe attached to the protopodite, and which bears thick paddle-shaped lamellae with short marginal setae (06). Key differences between the exopodite in these taxa include the shape of the lobe – subtriangular in *P. daziensis* versus semiovate in *R. abnormalis*, the presence of only up to nine lamellae in *P. daziensis* (compared to over a dozen in *R. abnormalis*), and the fact that *R. abnormalis* paddle-shaped lamellae are more elongate and more tightly imbricated. A comparable exopodite organization has also been recently confirmed in *Naraoia compacta* from the Burgess Shale (48), and tentatively suggested for *Emeraldella brocki* (37) and *Kuamaia lata* (06). Although *N. compacta* also features paddle-shaped lamellae with short marginal spines, it differs from *P. daziensis* and *R. abnormalis* in that the exopodite morphology consists of an elongate shaft, rather than a single robust lobe (Fig. 2c). The fifth to fourteenth post-antennal biramous appendages of *P. daziensis* share a similar overall morphology to that of the fourth pair, excluding their progressively smaller size posteriorly, but differ substantially in the morphology of the distal endopodite. The differentiation of a modified terminal claw and pre-terminal endopodite podomere into a robust moveable finger and fixed thumb respectively are not known in any other artiopod, nor the biramous appendages of any Cambrian euarthropod more broadly, as the endopodite termination usually consists of a single terminal claw that may carry smaller accessory spines that form a small functional foot (e.g. 02, 13, 37). In this context, the sub-chelate endopodites of the trunk appendages are morphologically more comparable with the walking legs in *L. polyphemus* (42), even if the terminal regions of the endopodites in *P. daziensis* are not fully developed into chelae as in horseshoe crabs. Subchelate limbs are also known from more phylogenetically distant euarthropods, such as pycnogonids and crustaceans (45), and are particularly well developed among burrowing representatives (49).

The presence of four morphologically distinct sets of appendages in *Pygmaclypeatus daziensis* (i.e. antennae, first to third post-antennal, fourth post-antennal, and fifth to fourteenth post-antennal) reflects some of the highest degrees of limb heteronomy expressed in any artiopod described to date. Although artiopods have been traditionally regarded as having largely homonomous series of biramous post-antennal appendages (06, 12, 13, 50), recent investigations record a growing number of cases of limb differentiation along the body. For example, the trilobite *Redlichia rex* from the Stage 4 Emu Bay Shale shows differences between the proportions of the anterior and posterior exopodites, even if their respective endopodites are morphologically similar (51). Limb heteronomy is more frequent among non-trilobite artiopods. The Miaolingian vicissicaudate *Emeraldella* features long antenniform antennae, one set of seemingly uniramous post-antennal appendages, biramous trunk limbs with some variability in terms of the relative length of the distal endopodite podomeres, and a pair of caudal flaps that likely represent modified exopodites (37, 52). *Sidneyia inexpectans* from the Wuliuan Burgess Shale also shows some differences between the (possibly uniramous) anterior and posterior post-antennal biramous trunk appendages (02), which might be linked with its durophagous feeding preferences (15, 16, 17, 53). However, the most notable cases of limb differentiation in Cambrian artiopods are found in the trilobitomorphs *Sinoburius lunaris* (22) and *Naraoia spinosa* (23) from Chengjiang, both of which have been recently redescribed based on micro-CT imaging. In addition to greatly reduced antennae, *S. lunaris* shows a striking differentiation in exopodite structure along the body: the first two pairs of post-antennal appendages have elongate stenopodous exopodites, whereas the remaining limb pairs have a more conventional exopodite shaft with lamellae. *Sinoburius lunaris* also has well-developed ridge-like endites on the trunk protopodites, whereas these are absent from the cephalic region (22). *Naraoia spinosa* also shows differentiation between the protopodite morphology in the cephalic and trunk regions, expressed as long spinose endites versus short blunt endites respectively, as well as more pronounced changes during ontogeny that suggest a shift from plankton/detritus feeding to scavenging (23). *Pygmaclypeatus daziensis* reflects possibly the highest degree of appendage heteronomy and functional specialization within Artiopoda described to date, given that differentiation is expressed in all components of the biramous limbs (i.e. protopodite, endopodite, exopodite), as well as between the cephalic and trunk regions (Fig. 3).

### Functional morphology and paleoecology

The new data on the ventral organization of *Pygmaclypeatus daziensis* prompt a reconsideration of its mode of life and paleoecology. Whereas earlier studies suggested an epibenthic habitus based only on the dorsoventrally flattened exoskeleton, the presence of well-developed paddle-like lamellae on the trunk exopodites suggests a considerable swimming prowess. Similar to *Naraoia compacta* (48), the exopodites would have likely produced effective propulsion during the power-stroke of a typical metachronal wave, and allowed for low drag during the recovery phase due to the pivot-like articulation of the individual paddles relative to the exopodite lobe. Taken together, the imbrication of the trunk exopodites coupled with the small overall body size (below 2 cm) suggest that *P. daziensis* would have been able to swim in the water column periodically, in line with an active nektobenthic mode of life. Based on recent studies supporting a gill-like function for the trilobite exopodite (54), the well-developed exopodites of *P. daziensis* may have effectively conveyed a large surface area for gas exchange thanks to the presence of more than 200 individual paddle-shaped lamellae (Fig. 2, 3).

The functional morphology of the protopodite and endopodites further informs the feeding habits of *P. daziensis*. The presence of delicate spinose endites on the protopodite of most biramous appendages suggests a diet consisting of soft-food items and organic-rich particles, compatible with scavenging/detritus feeding. The morphology of the endites rules out a durophagous diet, as shell crushing requires robust molariform gnathobases on transversely elongate protopodites, as seen in *Sidneyia inexpectans, Redlichia rex* and *Limulus polyphemus* (17). The absence of endites on the ventral side of the endopodites suggests that food processing mainly took place proximally within the trunk protopodites that form a food groove leading to the posteriorly-directed mouth opening underneath the hypostome (Fig. 2d), whereas the cephalic endopodites would be better suited for object manipulation in front of the body. The distal portion of all the endopodites would have been primarily used for locomotion. In particular, the strong ventral curvature of the legs, coupled with the presence of subchelate terminations on the trunk region consisting of the fixed thumb and the moveable finger appear to be well suited for a more specialized grabbing/anchoring function on hard substrates, and possibly burrowing behavior, as observed in extant forms such as amphipods (54, 55, 56). The similarities between *P. daziensis* and *L. polyphemus* (see previous section) suggest that these organisms shared a broadly comparable mode of life, the only important exception being that the lack of robust gnathobases in the former argues against effective durophagy as observed in *L. polyphemus* (17, 53), and instead favors a diet based on soft food items and/or decaying organic matter.

### Implications for artiopod evolution

The detailed three-dimensional appendage organization of *Pygmaclypeatus daziensis* contributes towards a better understanding of the paleobiology of this taxon (Fig. 3), and also has direct implications for reconstructing the evolutionary history of Artiopoda. The results of our phylogenetic analyses consistently resolve *P. daziensis* as the sister-taxon to *Retifacies abnormalis*, supported by the organization of the exopodite (single lobe with paddle-shaped lamellae with marginal setae), and the presence of both a pygidium and a multiarticulated tailspine; these results concur with earlier analyses that also support a close relationship between these two taxa (10, 11, 29). Implied weights (*k*=4) support the position of *P. daziensis* + *R. abnormalis* as the earliest branching clade within Trilobitomorpha (Fig. 4b), although its precise placement relative to the former is unresolved under equal weights (Fig. 4a). Despite the difference in these topologies, both analyses support an early-branching position within Artiopoda, which allows reconstructing the ancestral organization for these euarthropods.

The morphology of *P. daziensis* and *R. abnormalis* includes some characters that are widespread – and possibly ancestral – for trilobitomorphs, such as the conterminant hypostome and the presence of a pygidium. The presence of a multiarticulated tailspine appears to be unique to these two taxa within Trilobitomorpha, even though this character is known from more distantly related Cambrian forms such as *Molaria spinifera* (58); the latter case is most likely a result of homoplasy. A potentially significant new insight consists of the clarification of the detailed exopodite structure in *P. daziensis* and *R. abnormalis*, and its bearing for understanding the evolution of this structure among Trilobitomorpha. The exopodite of *P. daziensis* and *R. abnormalis* is unique as it consists of a single lobe with paddle-shaped lamellae, whereas the exopodite of most other trilobitomorphs is differentiated into a proximal lobe with lamellae and a distal lobe with short setae. As highlighted in recent work (48), and further demonstrated in the present study, the repeated occurrence of paddle-shaped lamellae in artiopods could argue in favour of an ancestral organization of the exopodites, which has not been fully appreciated due to the delicate nature of these structures, and the bias caused by compaction during burial. We explored this hypothesis through the optimization of paddle-shaped lamellae (Char. 15, state 2; see Supplementary Information) using the character mapping function in TNT (34, 59). Under equal weights (Fig. 4a), the ancestral organization of the exopodite lamellae for Artiopoda remains uncertain in all four most parsimonious trees recovered due to topology conflict resulting in a polytomy at the base of this clade. By contrast, the two most parsimonious trees recovered under implied weights (*k*=4) support the paddle-shaped lamellae as ancestral for Artiopoda in line with our hypothesis (Fig. 4b). A more definitive test to this hypothesis will require further investigations on the detailed appendage morphology of several early-branching artiopods (e.g. *Squamacula*, Protosutura; see 57), trilobitomorphs (e.g. Nektaspida, Xandarellida, Conciliterga; see 01, 6, 7) and vicissicaudates (e.g. Xenopoda, Cheloniellida, Aglaspidida; see 3, 61).

The presence of only five endopodite podomeres (including the terminal claw) in *P. daziensis* is remarkable in the context of Artiopoda, as most of these organisms feature the typical seven-segmented endopodite that is considered as ancestral for crown-group Euarthropoda in a broad sense (45, 47). Evidence for endopodites with less than seven podomeres in artiopods is rare. Possible cases include the reduced endopodites on the anteriormost biramous limbs of *Sinoburius lunaris* (22) and *Emeraldella* (37, 52) based on their small size compared with trunk appendages, but in both cases, the proximal portions of the endopodites are not well known. *P. daziensis* represents the only Cambrian artiopod for which all the post-antennal biramous appendages have a five-segmented endopod, as even its sister taxon *Retifacies abnormalis* shows more conventional limbs (06). The Devonian cheloniellid *Cheloniellon calmani* (62) is the only other known artiopod with consistently five-segmented endopodites throughout the body confirmed to date, although their structure is much closer to a generalized walking leg when compared to *P. daziensis*. Beyond Artiopoda, the five-segmented endopodite is widespread among mandibulates, and in particular pancrustaceans, whereas the seven-segmented endopodite is common among chelicerates (45, 47). Although the number of endopodite podomeres carries little phylogenetic information by itself, the biramous appendage construction of *P. daziensis* can be confidently interpreted as a derived trait among artiopods, with rare instances of convergence as observed in the phylogenetically and biostratigraphically distant *C. calmani* (62).

Another significant implication of this study is that it challenges the traditional notion that trilobitomorphs, and more broadly artiopods, are characterized by largely homonomous post-antennal appendage pairs (01, 06, 13, 63). Instead, *P. daziensis* reveals that some of the earliest branching representatives of this clade already demonstrate substantial limb differentiation (Fig. 4). Although *P. daziensis* is the latest example, it is by far not the only one, as variable degrees of post-antennal appendage specialization have been described in the literature (see discussion above). Most of the main artiopod clades include at least one representative for which there is substantial evidence of limb differentiation, including also Nektaspida (e.g. *Naraoia spinosa* (23)), Xandarellida (e.g. *Sinoburius lunaris* (22)), Trilobita (e.g. *Redlichia rex* (51)), and Vicissicaudata (e.g. *Emeraldella* (37, 52), *Sidneyia inexpectans* (02)). It is notable that these taxa are among the most complete and recently redescribed representatives of their respective groups, which suggests that anteroposterior limb differentiation might be more widespread within Artiopoda than it is currently considered. Other clades, such as Conciliterga, include taxa that have not been revised in at least the last decade, or that contain forms mostly known from the dorsal exoskeleton without complete appendage series. In this context, it appears likely that the archetypical homonomous appendage series of trilobites could represent a phylogenetically derived trait, rather than ancestral as commonly considered historically (e.g. 13, 45, 50, 63). Further morphological investigations on the appendage structure of Artiopoda that provide more comparable information between species are required, but the increased availability of new data on Chengjiang euarthropods through micro-CT imaging offers great promise for testing this hypothesis.

## ACKNOWLEDGMENTS

This study was supported by the NSFC grant (41861134032), DFG grant (Me-2683/10-1), Natural Science Foundation of Yunnan Province grants (2015HA021, 2018FA025, 2018IA073 and 2019DG050), and the Harvard China Fund.

## AUTHOR CONTRIBUTIONS

M.S., R.R.M., J.O.-H. and Y.L. designed the research. X.H., R.R.M., J.O.-H., and Y.L. secured the funding. X.H. and D.Z. collected all the material. R.R.M. and H.M. scanned YKLP 13929 and YKLP 11428, respectively). X.C. documented all the specimens with light photography. M.S. and Y.L. processed the tomographic models. M.S. rendered the 3D models. J.O.-H. performed the phylogenetic analysis. J.B. helped with the supplementary figure. M.S., J.O.-H. and R.R.M. wrote the paper with input from all the other coauthors. All authors participated in the interpretation of the material and the discussions.

## SUPPLEMENTARY FIGURES

**Extended Data Figure 1.**
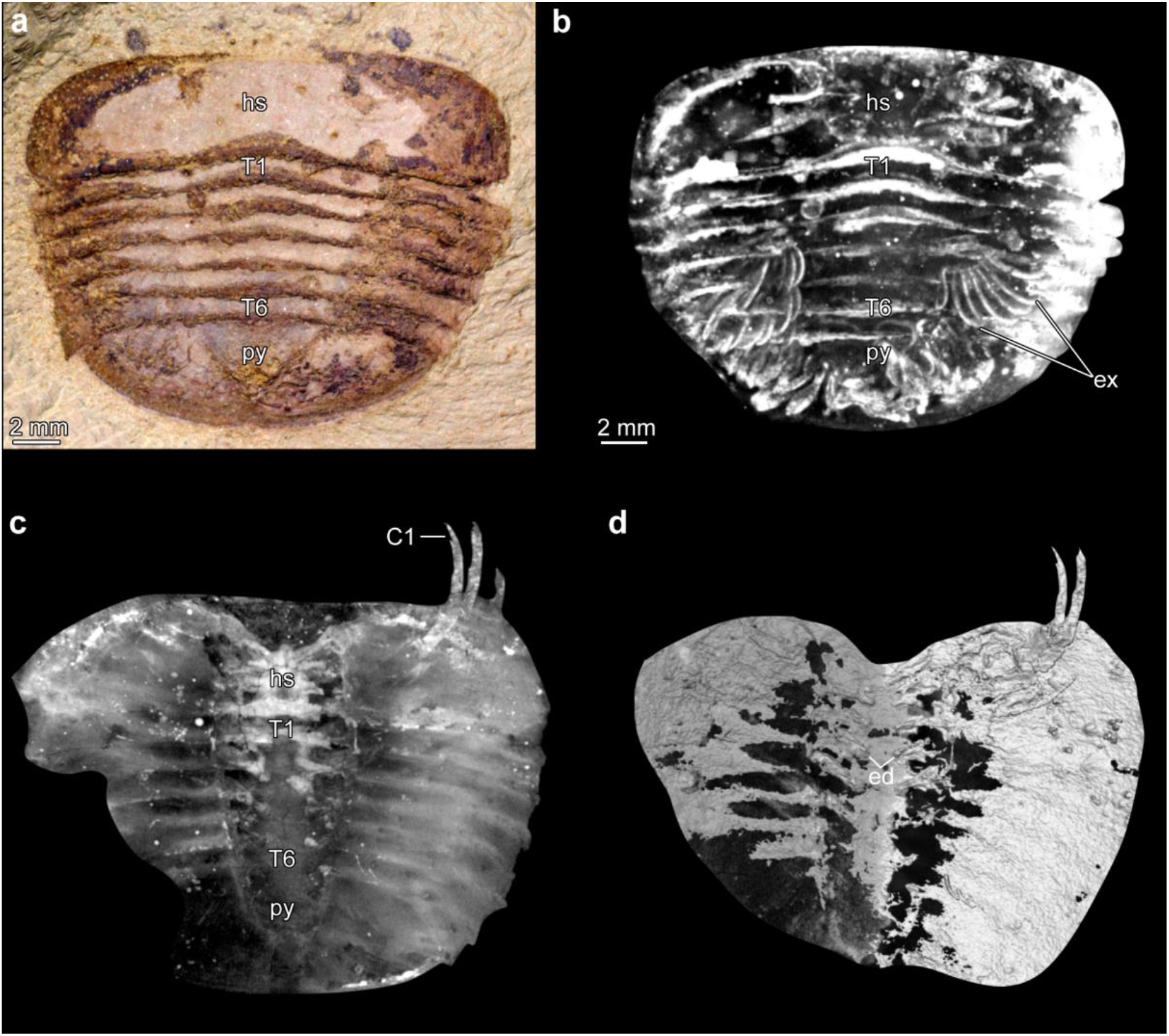
Additional specimens of *Pygmaclypeatus daziensis*. **a**. Specimen YKLP 13929, photographed under light microscopy. **b**. Dorsal view of three-dimensional computer model. **c**. Specimen YKLP 13928, volume rendering. **d**. Isosurface model showing ventral endites. Abbreviations: *Cn*, cephalic post-antennal appendage pair number; *ed*, endites; *ex*, exopodite; *Tn*, tergite number.

## REFERENCES

1. Edgecombe, G.D. and Ramsköld, L., 1999. Relationships of Cambrian Arachnata and the systematic position of Trilobita. Journal of Paleontology, 73 (2): 263–287.

2. Stein, M., 2013. Cephalic and appendage morphology of the Cambrian arthropod Sidneyia inexpectans. Zoologischer Anzeiger-A Journal of Comparative Zoology, 253(2), pp. 164–178.

3. Ortega-Hernández, J., Legg, D.A. and Braddy, S.J., 2013. The phylogeny of aglaspidid arthropods and the internal relationships within Artiopoda. Cladistics, 29(1), pp. 15–45.

4. Bond, A.D. and Edgecombe, G.D., 2021. Phylogenetic response of naraoiid arthropods to early–middle Cambrian environmental change. Palaeontology, 64(1), pp. 161–177.

5. Wilmot, N.V., 1990. Primary and diagenetic microstructures in trilobite exoskeletons. Historical Biology, 4(1), pp. 51–65.

6. Hou X. G, Bergström J. 1997. Arthropods of the Lower Cambrian Chengjiang fauna, Southwest China. Fossils Strata. 45: 1–116.

7. Mayers, B., Aria, C. and Caron, J.B., 2019. Three new naraoiid species from the Burgess Shale, with a morphometric and phylogenetic reinvestigation of Naraoiidae. Palaeontology, 62(1), pp. 19–50.

8. Budd, G.E., 2011. Campanamuta mantonae gen. et. sp. nov., an exceptionally preserved arthropod from the Sirius Passet Fauna (Buen Formation, lower Cambrian, North Greenland). Journal of Systematic Palaeontology, 9(2), pp. 217–260.

9. Stein, M., Budd, G.E., Peel, J.S. and Harper, D.A., 2013. Arthroaspis n. gen., a common element of the Sirius Passet Lagerstätte (Cambrian, North Greenland), sheds light on trilobite ancestry. BMC Evolutionary Biology, 13(1), pp. 1–34.

10. Paterson, J.R., Edgecombe, G.D., García-Bellido, D.C., Jago, J.B. and Gehling, J.G., 2010. Nektaspid arthropods from the lower Cambrian Emu Bay Shale Lagerstätte, South Australia, with a reassessment of lamellipedian relationships. Palaeontology, 53(2), pp. 377–402.

11. Paterson, J.R., García-Bellido, D.C. and Edgecombe, G.D., 2012. New artiopodan arthropods from the early Cambrian Emu Bay Shale Konservat-Lagerstätte of South Australia. Journal of Paleontology, 86(2), pp. 340–357.

12. Wills, M.A., Briggs, D.E.G. and Fortey, R.A., 1998. Evolutionary correlates of arthropod tagmosis: scrambled legs. In Arthropod relationships (pp. 57–65). Springer, Dordrecht.

13. Zeng, H., Zhao, F., Yin, Z. and Zhu, M., 2017. Appendages of an early Cambrian metadoxidid trilobite from Yunnan, SW China support mandibulate affinities of trilobites and artiopods. Geological Magazine, 154(6), pp. 1306–1328.

14. Vannier, J. and Chen, J.Y., 2002. Digestive system and feeding mode in Cambrian naraoiid arthropods. Lethaia, 35(2), pp. 107–120.

15. Zacaï, A., Vannier, J. and Lerosey-Aubril, R., 2016. Reconstructing the diet of a 505-million-year-old arthropod: Sidneyia inexpectans from the Burgess Shale fauna. Arthropod Structure & Development, 45(2), pp. 200–220.

16. Bicknell, R.D., Paterson, J.R., Caron, J.B. and Skovsted, C.B., 2018. The gnathobasic spine microstructure of recent and Silurian chelicerates and the Cambrian artiopodan Sidneyia: functional and evolutionary implications. Arthropod Structure & Development, 47(1), pp. 12–24.

17. Bicknell, R.D., Holmes, J.D., Edgecombe, G.D., Losso, S.R., Ortega-Hernández, J., Wroe, S. and Paterson, J.R., 2021. Biomechanical analyses of Cambrian euarthropod limbs reveal their effectiveness in mastication and durophagy. Proceedings of the Royal Society B, 288(1943), p. 20202075.

18. Zhai, D., Ortega-Hernández, J., Wolfe, J.M., Hou, X., Cao, C. and Liu, Y., 2019. Three-dimensionally preserved appendages in an early Cambrian stem-group pancrustacean. Current Biology, 29(1), pp. 171–177.

19. Zhai, D., Williams, M., Siveter, D.J., Harvey, T.H., Sansom, R.S., Gabbott, S.E., Siveter, D.J., Ma, X., Zhou, R., Liu, Y. and Hou, X., 2019. Variation in appendages in early Cambrian bradoriids reveals a wide range of body plans in stem-euarthropods. Communications biology, 2(1), pp. 1–6.

20. Liu, Y., Ortega-Hernández, J., Chen, H., Mai, H., Zhai, D. and Hou, X., 2020. Computed tomography sheds new light on the affinities of the enigmatic euarthropod Jianshania furcatus from the early Cambrian Chengjiang biota. BMC Evolutionary Biology, 20, pp. 1–17.

21. Liu, Y., Ortega-Hernández, J., Zhai, D. and Hou, X., 2020. A reduced labrum in a Cambrian great-appendage Euarthropod. Current Biology, 30(15), pp. 3057–3061.

22. Chen, X., Ortega-Hernández, J., Wolfe, J.M., Zhai, D., Hou, X., Chen, A., Mai, H. and Liu, Y., 2019. The appendicular morphology of Sinoburius lunaris and the evolution of the artiopodan clade Xandarellida (Euarthropoda, early Cambrian) from South China. BMC evolutionary biology, 19(1), pp. 1–20.

23. Zhai, D., Edgecombe, G.D., Bond, A.D., Mai, H., Hou, X. and Liu, Y., 2019. Fine-scale appendage structure of the Cambrian trilobitomorph Naraoia spinosa and its ontogenetic and ecological implications. Proceedings of the Royal Society B, 286(1916), p. 20192371.

24. Zhang, X.L., Shu, D.G. and Erwin, D.H., 2007. Cambrian naraoiids (Arthropoda): morphology, ontogeny, systematics, and evolutionary relationships. Journal of Paleontology, 81(sp68), pp. 1–52.

25. Størmer, L. 1944. On the relationships and phylogeny of fossil and recent Arachnomorpha. Skrifter Utgitt av Det Norske Videnskaps-Akademi i Oslo. I. Matematisk-Naturvitenskapelig Klasse 1944 5: 158 pp.

26. Zhang, X., Han, J. and Shu, D., 2000. A new arthropod Pygmaclypeatus daziensis from the early Cambrian Chengjiang Lagerstätte, South China. Journal of Paleontology, 74(5), pp. 979–983.

27. Xu, G. H. 2004. New specimens of rare arthropods from the early Cambrian Chengjiang fauna, Yunnan, China. Acta Palaeontologica Sinica 43(3): 325–331.

28. Hou, X.G., Siveter, D.J., Siveter, D.J., Aldridge, R.J., Pei-Yun, C., Gabbott, S.E., Xiao-Ya, M., Purnell, M.A. and Williams, M., 2017. The Cambrian fossils of Chengjiang, China: The flowering of early animal life. John Wiley & Sons.

29. Legg, D.A., Sutton, M.D. and Edgecombe, G.D., 2013. Arthropod fossil data increase congruence of morphological and molecular phylogenies. Nature communications, 4(1), pp. 1–7.

30. Gabbott, S.E., Xian-Guang, H., Norry, M.J. and Siveter, D.J., 2004. Preservation of Early Cambrian animals of the Chengjiang biota. Geology, 32(10), pp. 901–904.

31. Zhu, M., Babcock, L.E. and Steiner, M., 2005. Fossilization modes in the Chengjiang Lagerstätte (Cambrian of China): testing the roles of organic preservation and diagenetic alteration in exceptional preservation. Palaeogeography, Palaeoclimatology, Palaeoecology, 220(1-2), pp. 31–46.

32. Limaye, A., 2012, October. Drishti: a volume exploration and presentation tool. In Developments in X-ray Tomography VIII (Vol. 8506, p. 85060X). International Society for Optics and Photonics.

33. Garwood, R. and Dunlop, J., 2014. The walking dead: Blender as a tool for paleontologists with a case study on extinct arachnids. Journal of Paleontology, 88(4), pp. 735–746.

34. Goloboff, P.A. and Catalano, S.A., 2016. TNT version 1.5, including a full implementation of phylogenetic morphometrics. Cladistics, 32(3), pp. 221–238.

35. Lankester ER. The structure and classification of Arthropoda. Quarterly J. Microscop. Sci. 1904;47: 523–582.

36. Zhang, X., Han, J., Zhang, Z., Liu, H. and Shu, D., 2004. Redescription of the Chengjiang arthropod Squamacula clypeata Hou and Bergström, from the Lower Cambrian, south-west China. Palaeontology, 47(3), pp. 605–617.

37. Stein, M. and Selden, P.A., 2012. A restudy of the Burgess Shale (Cambrian) arthropod Emeraldella brocki and reassessment of its affinities. Journal of Systematic Palaeontology, 10(2), pp. 361–383.

38. Ortega-Hernández, J., 2015. Homology of head sclerites in Burgess Shale euarthropods. Current Biology, 25(12), pp. 1625–1631.

39. Ortega-Hernández, J., Janssen, R. and Budd, G.E., 2017. Origin and evolution of the panarthropod head–a palaeobiological and developmental perspective. Arthropod structure & development, 46(3), pp. 354–379.

40. Zeng, H., Zhao, F., Niu, K., Zhu, M. and Huang, D., 2020. An early Cambrian euarthropod with radiodont-like raptorial appendages. Nature, 588(7836), pp. 101–105.

41. Whittington, H.B., 1980. Exoskeleton, moult stage, appendage morphology, and habits of the Middle Cambrian trilobite Olenoides serratus. Palaeontology 23: 171–204.

42. Haug, C. and Rötzer, M.A., 2018. The ontogeny of Limulus polyphemus (Xiphosura s. str., Euchelicerata) revised: looking “under the skin”. Development genes and evolution, 228(1), pp. 49–61.

43. Legg, D.A., 2014. Sanctacaris uncata: the oldest chelicerate (Arthropoda). Naturwissenschaften, 101(12), pp. 1065–1073.

44. Bruton, D.L., 1981. The arthropod Sidneyia inexpectans, Middle Cambrian, Burgess Shale, British Columbia. Philosophical Transactions of the Royal Society of London. B, Biological Sciences, 295(1079), pp. 619–653.

45. Boxshall, G.A., 2004. The evolution of arthropod limbs. Biological Reviews, 79(2), pp. 253–300.

46. Wolff, C. and Scholtz, G., 2008. The clonal composition of biramous and uniramous arthropod limbs. Proceedings of the Royal Society B: Biological Sciences, 275(1638), pp. 1023–1028.

47. Bruce, H.S., 2021. How to align arthropod legs. bioRxiv. (https://doi.org/10.1101/2021.01.20.427514)

48. Haug, C. and Haug, J.T., 2016. New insights into the appendage morphology of the Cambrian trilobite-like arthropod Naraoia compacta. Bulletin of Geosciences, 91(2), pp. 221–227.

49. Faulkes, Z., 2013. Morphological adaptations for digging and burrowing. Functional morphology and diversity, pp. 276–295.

50. Whittington, H.B. and Almond, J.E., 1987. Appendages and habits of the Upper Ordovician trilobite Triarthrus eatoni. Philosophical Transactions of the Royal Society of London. B, Biological Sciences, 317(1182), pp. 1–46.

51. Holmes, J.D., Paterson, J.R. and García-Bellido, D.C., 2020. The trilobite Redlichia from the lower Cambrian Emu Bay Shale Konservat-Lagerstätte of South Australia: systematics, ontogeny and soft-part anatomy. Journal of Systematic Palaeontology, 18(4), pp. 295–334.

52. Lerosey-Aubril, R. and Ortega-Hernández, J., 2019. Appendicular anatomy of the artiopod Emeraldella brutoni from the middle Cambrian (Drumian) of western Utah. PeerJ, 7, p. e7945.

53. Bicknell, R.D., Ledogar, J.A., Wroe, S., Gutzler, B.C., Watson III, W.H. and Paterson, J.R., 2018. Computational biomechanical analyses demonstrate similar shell-crushing abilities in modern and ancient arthropods. Proceedings of the Royal Society B, 285(1889), p. 20181935.

54. Vader, W., 1983. Prehensile pereopods in gammaridean Amphipoda. Sarsia, 68(2), pp. 139–148.

55. Holmquist, J.G., 1982. The functional morphology of gnathopods: importance in grooming, and variation with regard to habitat, in talitroidean amphipods. Journal of Crustacean Biology, 2(2), pp. 159–179.

56. Bousfield, E.L., 1965. Haustoriidae of New England (Crustacea: Amphipoda). Proceedings of the United States National Museum.

57. Hou, J.B., Hughes, N.C. and Hopkins, M.J., 2021. The trilobite upper limb branch is a well-developed gill. Science Advances, 7(14), p. eabe7377.

58. Whittington, H.B., 1981. Rare arthropods from the Burgess Shale, Middle Cambrian, British Columbia. Philosophical Transactions of the Royal Society of London. B, Biological Sciences, 292(1060), pp. 329–357.

59. Goloboff, P.A., Farris, J.S. and Nixon, K.C., 2008. TNT, a free program for phylogenetic analysis. Cladistics, 24(5), pp. 774–786.

60. Du, K.S., Ortega-Hernández, J., Yang, J. and Zhang, X.G., 2019. A soft-bodied euarthropod from the early Cambrian Xiaoshiba Lagerstätte of China supports a new clade of basal artiopodans with dorsal ecdysial sutures. Cladistics, 35(3), pp. 269–281.

61. Lerosey-Aubril, R., Zhu, X. and Ortega-Hernández, J., 2017. The Vicissicaudata revisited–insights from a new aglaspidid arthropod with caudal appendages from the Furongian of China. Scientific Reports, 7(1), pp. 1–18.

62. Stürmer, W. and Bergström, J., 1978. The arthropod Cheloniellon from the Devonian Hunsrück shale. Paläontologische Zeitschrift, 52(1), pp. 57–81.

63. Hughes, N.C., 2007. The evolution of trilobite body patterning. Annu. Rev. Earth Planet. Sci., 35, pp. 401–434.

