## Supplementary material for "Before trilobite legs: *Pygmaclypeatus daziensis* reconsidered and the ancestral appendicular organization of Cambrian artiopods": Pygmaclypeatus_V1_character_list.txt

    Phenotypic characters (numbered in Nexus format)

    Array characters
  1. nature of first appendage: raptorial appendage (0); antennae (1).
  2. number of podomeres on raptorial limb: four (0); three (1); two (2).
  3. raptorial appendages with flagellum: absent (0); present (1).
  4. length of distal spines on podomeres: absent or shorter than podomere (0); subequal to length of podomeres (1); longer than entire podomere series (2).
  5. number of post-ocular head segments: none (0); one (1); two (2); three (3); four (4); five (5); six (6); seven (7).
  6. nature of second appendage pair: biramous walking leg or antennae (0); uniramous walking leg (1).
  7. nature of third appendage pair: biramous walking leg (0); uniramous walking leg (1).
  8. appendages under cephalo-thoracic articulation: absent (0); present (1).
  9. trunk endopods: absent or reduced (0); present (1).
  10. trunk exopod structure (UPDATED): simple oval flap (0); exopod differentiated into proximal and distal lobes, the proximal lobe bearing lamellar setae, the distal lobe bearing short setae (1); numerous podomeres, each bearing a single seta (2); undivided lobe with lamellae (3); book gills (4).
  11. proximal exopod lobe: flattened lobe (0); slender shaft (1).
  12. distal exopod lobe: small to moderate sized flap (0); large, teardrop shaped lobe with long attachment (1).
  13. exopod with articulations: absent (0); present (1).
  14. imbricate exopod lamellae: absent (0); present (1).
  15. lamellae structure (UPDATED): thick, flat lamellae (0); delicate, comb-like lamellae (1); paddle-shaped lamellae with marginal setae (2).
  16. non-overlapping marginal setae on distal lobe: absent (0); small setae (1); long spines or filaments (2).
  17. gnathobasic limbs: absent (0); present (1).
  18. lateral eyes: absent (0); present (1).
  19. nature of lateral eyes: stalked (0); sessile (1).
  20. ventral eyes in front of head: absent (0); present (1).
  21. calcified eyes: absent (0); present (1).
  22. dorsal exoskeletal bulge: absent (0); present (1).
  23. Eye slits: absent (0); present (1).
  24. dorsal median eyes: absent (0); present (1).
  25. free head shield: absent (0); present (1).
  26. bivalved carapace: absent (0); present (1).
  27. cephalic doublure: absent (0); present (1).
  28. cephalon notched: absent (0); present (1).
  29. hypostome: median extension of the doublure, with no suture (0); natant, sclerite not in contact with doublure (1); with narrow overlap with pre-hypostomal sclerite (2); narrow attachment to doublure at hypostomal suture (3); absent (4).
  30. anterior sclerite: absent (0); present (1).
  31. ecdysial sutures: absent (0); present (1).
  32. position of ecdysial sutures: marginal (0); dorsal (1).
  33. elevated marginal rim: absent (0); present (1).
  34. marginal rim ornamented: absent (0); present (1).
  35. differentiated glabellar region: absent (0); present (1).
  36. head shield outline: genal spines (0); acute genal angles (1); rounded genal angles (2); lateral spine-like extensions of the carapace (3).
  37. posterior cephalic band: absent (0); present (1).
  38. pleural cephalic furrows: absent (0); present (1).
  39. cuticle sculpture: absent (0); present (1).
  40. mineralized cuticle: absent (0); present (1); State 2 (2).
  41. expanded lateral pleurae: absent (0); present (1).
  42. free thoracic tergites: absent (0); present (1).
  43. decopuling of tergites and limb pairs: absent (0); present (1).
  44. tergite articulations: tergites non-overlapping (0); extensive overlap of tergites (1); edge to edge pleural articulations (2).
  45. dorsal trunk effacement: trunk with defined (separate or fused) tergite boundaries (0); trunk tergite boundaries effaced laterally (1); trunk tergite boundaries completely effaced (2).
  46. cephalic articulation fused: absent (0); present (1).
  47. head shield extends over anterior tergites: overlap absent or identical to overlap between thoracic segments (0); head shield covers first thoracic tergite only (1); head shield covers multiple anterior trunk tergites (2).
  48. articulation with reduced tergite: absent (0); present (1).
  49. trunk narrowed anteriorly: absent (0); present (1).
  50. trunk tergites reflexed anteriorly: absent, boundaries transverse or reflexed posterolaterally (0); present (1).
  51. joints between posterior tergites functional: absent (0); present (1).
  52. posterior tergite with single axial spine: absent (0); present (1).
  53. radial pleurae: absent (0); present (1).
  54. raised axial region: absent or weakly defined (0); present and well defined (1).
  55. anterior tergal processes: absent (0); present (1).
  56. tuberculation of posterior tergite margin: absent (0); present (1).
  57. axial spines or nodes: absent (0); present (1).
  58. length of postabdomen: one segment (0); two segments (1); three segments (2); five segments (3).
  59. postabdomen without walking legs: absent (0); present (1).
  60. postabdomen as differentiated tergites: absent (0); present - tergite free (e.g. some aglaspidids, cheloniellids, Emeraldella, Sidneyia) (1); present - preterminal tergite fused with telson (most aglaspidids) (2).
  61. posterior tergites strongly curved: absent (0); present (1).
  62. pygidium: absent (0); present (1).
  63. position of anus: terminal, within telson (0); at base of telson (1).
  64. pygidium with median keel: absent (0); present (1).
  65. pygidium with broad median spine: absent (0); present (1).
  66. pygidium with lateral spines: absent (0); present (1).
  67. tailspine: absent (0); present (1).
  68. tailspine shape: spinose (0); paddle shaped (1); cap-like (2).
  69. length of tailspine: shorter than half the length of the trunk (0); longer than half the length of the trunk (1).
  70. marginal spines on tailspine: absent (0); present (1).
  71. tailspine with medial cleft or keel: absent (0); present (1).
  72. paddle with projections: absent (0); present (1).
  73. ventral sclerite covering anal region: absent (0); present (e.g. aglaspidids, Sidneyia) (1).
  74. paired modified appendages: absent (0); present (1).
  75. nature of preterminal appendages: uropods (0); furca (1).
  76. reduced carapace: absent (0); present (1).
  77. width of doublure: narrow to moderately wide (0); wide (1); covers entire ventral side of cephalon (2).
  78. length of thorax relative to pygidium: longer than pygidium (0); shorter than pygidium (1).
  79. articulating half rings: absent (0); present (1).
  80. axial furrows on trunk: absent (0); present (1).
  81. postventral plate medial attachment: narrow attachment by central portion (0); wide longitudinal attachment (1).
  82. dorsal eyes confined to anterior half: absent (e.g. Trilobita) (0); present (e.g. Aglaspidida) (1).
  83. dorsal eye abutting glabella anteriorly: absent (e.g. Trilobita) (0); present (e.g. Aglaspidida) (1).
  84. dorsal eyes merge anteriorly into cephalon: absent (e.g. Trilobita) (0); present and eyes are separate (e.g. Glypharthrus) (1); present and eyes are medially fused (e.g. Cyclopites) (2).
  85. nature of anal sclerite: anal plate (e.g. Sidneyia) (0); postventral plate (e.g. Aglaspidida) (1).
  86. nature of differentiated preterminal tergite: tergite with reduced pleurae (e.g. Weinbergina) (0); cylindrical tergite without pleurae (e.g. Aglaspidida, Cheloniellida, Emeraldella (1).
  87. pleural tips on posterior end with elongate spines: absent (0); present (1).
  88. antennal scale: absent (0); present (1).
  89. head shield with lateral notches: absent (0); present (1).
  90. tailspine with multiple articulations (NEW): absent (0); present (1). 
  91. shape of undivided expod lobe with lamellae (NEW): broad lobe (e.g. Retifacies, Pygmaclypeatus) (0); slender shaft (e.g. Naraoia compacta) (1). 
  92. head/trunk exopods heteronomous (NEW): absent, exopods with same morphology in head and trunk (0); present, exopods with different morphology in head and trunk (1). 
